# Fluorescence-based CRISPR interference system for controlled genetic repression and live single-cell imaging in mycobacteria

**DOI:** 10.1101/2024.10.05.616838

**Authors:** Janïs Laudouze, Vanessa Point, Wafaa Achache, Céline Crauste, Stéphane Canaan, Pierre Santucci

## Abstract

Mycobacterial genetics has played a pivotal role over the last 35 years in our understanding of mycobacterial physiology, pathogenesis and antibiotic resistance. Numerous approaches are now available worldwide to dissect the contribution of genes of interest in biological processes. However, many of these approaches can be fastidious, difficult to perform and time-consuming, especially when working with slow-growing mycobacteria or in bio-safety level two/three settings. The recent development of CRISPRi-mediated targeted gene repression has revolutionized the way research groups can perform genetics in mycobacteria, providing a fast, robust and efficient alternative to study the function of specific genes including essential genes. In this research letter, we report the development and validation of a new subset of fluorescence-based CRISPRi tools for our scientific community. The pJL series is directly derived from the original integrative pIRL2 and pIRL117 CRISPRi vectors and conserved all the elements required to perform inducible targeted gene repression. In addition, these vectors carry two distinct fluorescent markers for which the expression is driven by the strong and constitutive promotor *psmyc* to simplify the selection of recombinant clones. We demonstrate the functionality of these vectors by targeting the expression of the non-essential glycopeptidolipid translocase *mmpL4b* and the essential genes *rpoB* and *mmpL3*. Finally, we describe an efficient single-step procedure to co-transform mycobacterial species with this integrative genetic tool alongside replicative vectors. Such tools and approaches should be useful to foster discovery in mycobacterial research.

**Graphical Abstract:** 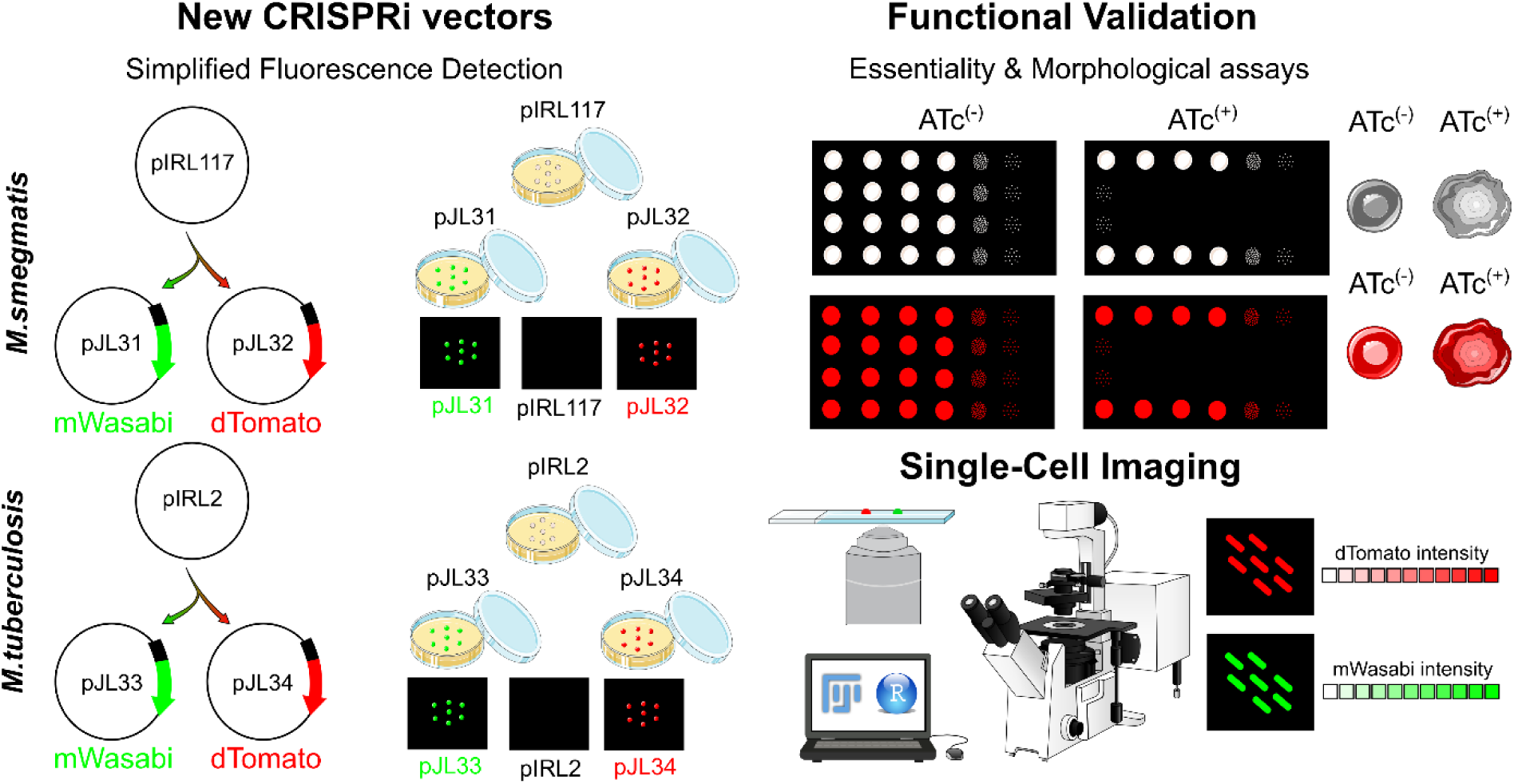

Development and validation of a new subset of E. coli-Mycobacteria shuttle vectors that enable simultaneous CRISPRI-mediated gene silencing and fluorescence based single-cell imaging.

## Introduction

The recent emergence and progress of CRISPR-based technologies has changed our way of thinking, designing and performing bacterial genetics, facilitating numerous aspects of genome editing. Among the technologies available, CRISPR-Cas-dependent genome modification has appeared to be successful in mycobacteria with studies reporting the generation of unmarked mutants in *Mycobacterium smegmatis* [1], *Mycobacterium marinum* [2], *Mycobacterium neoaurum* [3], *Mycobacterium abscessus* [4, 5] and *Mycobacterium tuberculosis* [2, 6].

As an alternative to CRISPR-Cas-mediated genome editing, the development of targeted gene silencing by CRISPR interference (CRISPRi) approaches has considerably revolutionized mycobacterial genetics [7–9]. Indeed, CRISPRi has shown numerous advantages in comparison to other genetic approaches such as its ability to target one or multiple (essential) genes alone or simultaneously, its inducible and therefore reversible nature alongside the fact that repression strength is tuneable. Altogether, this makes CRISPRi a very attractive and laboursaving system [10]. Moreover, the use of advanced whole-genome CRISPRi screen pioneered in the Rock research laboratory and further applied in different independent groups, has recently emerged as a unique and powerful methodology to perform functional genomics and further dissect mycobacterial physiology, pathogenesis and identify new drug targets [9, 11–13].

Another great advantage of targeted repression through CRISPRi in comparison to conventional two-step homologous recombineering strategies [14–16], relies on its single transformation/selection step [9, 10], hence leading to a faster generation of recombinant strains that can be subsequently validated before addressing the biological question(s) of interest. Unfortunately, despite this advantage, selection of recombinant clones remains fastidious in mycobacteria, with numerous species displaying very high-frequency of spontaneous mutations observed when selecting on conventional laboratory antibiotics such as hygromycin or kanamycin [4, 17]. Therefore, clonal selection, expansion and genetic validation of the newly generated recombinant strains can be time consuming. To overcome such limitation, additional selection markers conferring easily detectable phenotypes such as the beta-galactosidase *lacZ* or the catechol 2,3 dioxygenase *xylE* have been widely used in the past [15, 17]. More recently, genes encoding fluorescent proteins such as GFP, mWasabi or dTomato have been extensively used [18–21], thus facilitating generation and selection of bacterial mutant strains.

In addition to this selection step, genetically encoded fluorescent markers can be used as proxy or read out into a wide range of biological assays. In that context, quantitative fluorescence imaging has emerged as one of the most powerful technology in biological science with a wide range of applications in microbiology, non-exhaustively including drug susceptibility testing, intracellular replication or host/tissue colonization monitoring, gene-reporter studies or the investigation of protein subcellular localisation [22–27]. Seminal study by de Wet and colleagues, combining high-throughput inducible CRISPRi and image-based analyses have further demonstrated how coupling such technologies are critical to further understand mycobacterial physiology and potential drug mechanism of action [28].

In this research letter, we report the development of a new generation of four CRISPRi-based vectors that harbour the mWasabi or dTomato-coding sequence placed under the control of the strong constitutive *psmyc* promotor. Addition of these fluorescence expression cassettes facilitates the selection of recombinant clones and make possible their subsequent imaging by live single-cell fluorescence microscopy. Functional validation of such constructs was performed by targeting the expression of well-characterised mycobacterial genes such as the trehalose mono-mycolate translocase *mmpL3*, the RNA polymerase subunit-β *rpoB* and the glycopeptidolipid translocase *mmpL4b*. Finally, we describe a simple single-step procedure that allows co-transformation of CRISPRi-based vectors with episomal vectors encoding alternative fluorophores. Such methodology enables to rapidly isolate co-transformants in both fast and slow-growing bacteria, including the non-pathogenic environmental strain *M. smegmatis* and the tubercle bacilli, *M. tuberculosis*. These tools and approaches aim at facilitating complex investigations based on genetic alterations and dual fluorescence imaging in our scientific community.

## Material and Methods

### Mycobacterial strains and culture conditions

*Mycobacterium smegmatis* mc^2^ 155 and *Mycobacterium tuberculosis* ATCC 25117 H37Ra reference strains were used in this study. All mycobacterial strains were routinely grown in Middlebrook 7H9 broth (BD Difco, #271310) supplemented with 0.2% glycerol (Euromedex; #EU3550), 0.05% Tween 80 (Sigma-Aldrich; #P1754). In the case of *M. tuberculosis* 10% oleic acid, albumin, dextrose, catalase (OADC enrichment; BD Difco; #211886) was also added. All cultures were maintained at 37°C with shaking.

When required hygromycin B (Toku-E; #H007) or kanamycin (Euromedex; #UK0010D) were used as selection markers for transformants in Middlebrook 7H10 (BD Difco; #262710) or Middlebrook 7H11 (BD Difco; #283810) solid medium or for the culture maintenance of the fluorescent strains at a final concentration of 50 mg/L for *M. smegmatis* and *M. tuberculosis*.

### Cloning and generation of new fluorescent-based plasmids

All molecular biology experiments including plasmid selection, expansion and maintenance were performed in the reference strain *Escherichia coli* DH10B (Invitrogen). *E. coli* was grown into LB Broth (BD Difco; #244620) or in LB solid medium containing Agar Agar Bacteriological (Euromedex; #1330). When required hygromycin B (final concentration of 200 mg/L) or kanamycin (final concentration of 50 mg/L) were used as a selection marker for transformants on solid medium or for the culture maintenance of the recombinant *E. coli* strains. All plasmids and primers used in this study have been listed in supplementary information in Table S1 and Table S2. pJL29 (Addgene #227426) and pJL30 (Addgene #227427) are fluorescent-based vectors that harbour the *mWasabi* and *dTomato* genes under the control of the *psmyc* promotor. These vectors are derived from the pUV15-pHGFP, pTEC15 and pTEC27 vectors [25, 29]. Briefly, the *mWasabi* and *dTomato* genes were amplified by PCR (NEB; #M0491L) with the primers #P1-P2 and #P3-P4 by using the template pTEC15 and pTEC27, respectively (Addgene plasmids #30174 & #30182 kindly gifted by Lalita Ramakrishnan) [25]. DNA fragments were purified, digested with SphI and HindIII restriction enzymes (NEB; #R3182L and #R3104L). Simultaneously, pUV15-pHGFP (Addgene plasmid #70045, kindly gifted by Sabine Ehrt) [29] was digested by SphI and HindIII to remove the pH-GFP coding sequence. Purified and digested inserts and pUV15-pHGFP vector were then ligated using T4 DNA ligase (NEB; #M0202S) to generate the pJL29 and pJL30 vectors.

pJL31 (Addgene #227428), pJL32 (Addgene #227429), pJL33 (Addgene #227430) and pJL34 (Addgene #227431) are fluorescent-based CRISPRi vectors derived from the original pIRL117 and pIRL2 backbones respectively [11]. Both *mWasabi* and *dTomato* genes with their *psmyc* promotors were amplified by PCR using the primers #P5-P2 and #P5-P4 from pJL29 and pJL30 templates. DNA fragments were purified and ligated into HpaI-digested pIRL117 and pIRL2 (NEB; #R0105L) to generate the pJL31/pJL32 and pJL33/pJL34 vectors, respectively. All vectors generated were fully sequenced by whole plasmid sequencing (Eurofins Genomics).

### Molecular cloning of CRISPRi vectors

pIRL117 and pJL32 were used as proof of concept to perform CRISPRi-mediated silencing validation experiments. Cloning of the sgRNA targeting sequences was performed as previously described [10, 11]. Briefly, oligonucleotides were designed to target the non-template strand of the target gene ORF using the Peeble database (https://pebble.rockefeller.edu/) [10, 11]. Both top and bottom single-stranded DNA oligonucleotides (IDT) were pooled and annealed before being cloned into CRISPRi plasmid backbones. For cloning, CRISPRi plasmids were digested with BsmBI-v2 (NEB #R0739L) and further gel purified (Macherey-Nagel; #740609.50). For each individual single-guide ribonucleic acid (sgRNA) targeting sequence, two complementary annealed oligonucleotides with appropriate sticky end overhangs were ligated (NEB; #M0202M) into the desired BsmBI-v2 digested plasmid backbone. Successful cloning was confirmed by Sanger sequencing (Eurofins Genomics) using the primer #P6. The list of sgRNA targeting sequences #P7-P12 and generated plasmids used for constructing individual CRISPRi strains can be found in Table S1 and Table S2.

### Mycobacterial transformation and selection

Preparation of competent mycobacterial cells and their subsequent electroporation was performed as previously described [30]. Briefly, 100 mL of exponentially growing cells at an approximate optical density 600nm (OD_600nm_) of ∼0.4-0.8 were washed 5 times either at room temperature or at 4°C in sterile aqueous solution containing 10% glycerol and 0.05% Tyloxapol (Sigma; #T8761). At each step the volume washing solution was decreased (*e.g* 50 mL, 40 mL, 30 mL, 20 mL and 10 mL) and the final resuspension was performed in 1/50 to 1/100 of the original volume of culture. Then, approximately 500 ng-1 µg of vectors of interest were added to 100-200 µL of competent cells and placed into an electroporation cuvette of 0.2 mm gap. Electroporation settings were selected according to each reference strain [30]. For *M. tuberculosis* one single pulse of 2.5 kV, 25 μF, with the pulse-controller resistance set at 1,000 Ω resistance was performed whereas for *M. smegmatis* a single-used with 2.5 kV, 10 μF and 600 Ω was used.

Electroporated cells were transferred in 1-5 mL of complete 7H9 broth devoid of antibiotics and further incubated at 37°C for 4 h for *M. smegmatis* and 24 h for *M. tuberculosis* before plating. Transformants were selected on complete Middlebrook 7H10 agar containing the appropriate concentration of hygromycin B and/or kanamycin. Plates were incubated at 37°C for 4–7 days for *M. smegmatis* and 3-4 weeks for *M. tuberculosis*.

### Detection of mWasabi or dTomato expressing bacteria at the macro- and single-colony level using conventional bench-top imaging system

Approximately 10^6^ colony forming units (CFU) of exponentially growing *M. smegmatis* were used to inoculate conventional 7H10 plates. For single colonies visualisation, 10^6^ CFU were spotted and separated using the quadrant streak plating technique. For visualisation of macro-colonies, 10^6^ CFU were spotted as single 5-10 µL drop and allowed to dry for few minutes. In both cases, plates were incubated at 37°C, visual inspection of growth and scanning was performed after 2-4 days. For fluorescence detection in 7H9 broth, approximately 1-5×10^7^ CFU of recombinant bacteria harbouring the plasmids of interest were dispensed into standard flat-bottom 96 well-plates (Thermo-Fisher Scientific; #167008).

Low-resolution imaging of bacterial colonies, spots or liquid cultures was performed using the ChemiDoc^TM^ MP Imaging System and the Image Lab software version 6.1.0 (Bio-Rad). Bright light acquisition was performed by using the “White Epi Illumination” setting combined with the “Standard Filter”. Green fluorescence acquisition was performed by using the “Blue Epi Illumination” setting combined with the “530/28 nm Filter”. Finally, red fluorescence acquisition was performed by using the “Green Epi Illumination” setting combined with the “605/50 nm Filter”. When required individual raw images were exported from Image Lab as TIFF files 600 dpi and displayed with the open-source software Fiji/Imagej (https://imagej.net/software/fiji/) [31], to generate the final micrographs, including the merge channel.

### Assessment of colony morphology alterations on solid medium

For *M. smegmatis* morphotype assays, experiments were performed on conventional 7H10 plates containing 0.05% Tween-80 in the presence or absence of the inducer anhydrotetracycline (ATc) at a final concentration of 100 ng/mL [10]. When mentioned 100 µg/mL of Congo-Red was added to the media as previously described [32, 33]. Low-resolution imaging of colonies morphologies was performed described above using the ChemiDoc^TM^ MP Imaging System. Briefly, bright light acquisition was performed by using the “White Epi Illumination” setting combined with the “Standard Filter” whereas red fluorescence acquisition was performed by using the “Green Epi Illumination” setting combined with the “605/ 50nm Filter”. High-resolution observations of colony morphologies were performed using a stereomicroscope coupled with Canon EOS 550D camera (Canon). Images acquisition was performed using the manual focus and resolution of each image was 5184 x 3456 pixels. When required Individual raw images were exported as TIFF files and displayed with the open-source software Fiji/Imagej (https://imagej.net/software/fiji/) [31].

### Lipid extraction and semi-quantitative TLC analysis

Recombinant *M. smegmatis* strains harbouring pIRL117, pIRL117_*mmpL4b*, pJL32 and pJL32_*mmpL4b* were used to inoculate 200 mL 7H9 broth at initial OD_600nm_ of ∼0.025 in the presence or in the absence of 100 ng/mL of the ATc inducer. Cultures were grown simultaneously at 37°C under shaking at approximately 180-200 rpm. After 48 h of incubation, cultures were then centrifuged at 4,000 rpm for 15 min at 4°C and washed twice with distilled water. Pellets were frozen in liquid nitrogen and lyophilized overnight. Dry pellets were weighed to determine the exact mass of bacterial for normalization calculations before lipid extraction.

Total lipids were extracted as previously described [34] with slight modifications. Briefly, lipids from dry pellet were incubated for 16 h with CHCl_3_-CH_3_OH (1:2; *v/v*), at room temperature under shaking. Residual pellets were re-extracted for 16 h with the same solvents, but using alternative ratios (1:1 and 2:1 *v/v*). All the three organic phases were pooled and concentrated under reduced pressure. Samples were re-suspended in a CHCl_3_-CH_3_OH solution (3:1, *v/v*), and washed with 0.3% (*w/v*) NaCl in water. Organic and aqueous phases were separated by centrifugation 4,000 rpm for 10 min, and only the organic phases were conserved and dried over MgSO_4_. Samples were finally evaporated under a nitrogen stream, weighed, and re-suspended in a CHCl_3_-CH_3_OH solution (3:1, *v/v*). Equal volume of extract containing glycopeptidolipids (GPL) were deposited using a semi-automated sample application system LINOMAT 5 (CAMAG) and further separated on TLC (Silica Gel 60, Merck) by using CHCl_3_-CH_3_OH (90:10, *v/v*) as eluent. GPL were further visualized by vaporization of 0.2% (*v/v*) anthrone in a 20% sulfuric acid-ethanol solution. Finally, each resolved plate was heated at 120°C for 2-3 min using TLC Plate Heater III (CAMAG), scanned using a ChemiDoc^TM^ MP Imaging System, and densitometric analyses was performed using the built-in ImageLab^TM^ allowing to determine the relative content of each sample.

### Validation of targeted gene repression of essential genes using spot CFU-based assays

Assessment of genetic essentiality was performed by performing serial dilutions of *M. smegmatis* recombinant strains and plated them on 7H10 media in the presence or the absence of the ATc inducer (final concentration 100 ng/mL). Exponentially growing bacteria were normalised to an OD_600nm_ of 0.1 corresponding to approximately 1.10^7^ CFU per mL. Ten-fold serial dilutions were performed and 10 µL of each dilution (10^0^ to 10^-6^) was plated on each plate containing or not ATc. Plates were incubated at 37°C and visual inspection of growth was performed between day 2-4. Scanning of the plates was performed using the ChemiDoc^TM^ MP Imaging System as described above.

### Live-single cell imaging by fluorescence microscopy

All fluorescence microscopy experiments were performed with exponentially growing cells that were cultivated in 7H9 broth at an OD_600nm_ of ∼0.4-0.8. Briefly, 1 mL of cells were washed twice in sterile phosphate buffer saline (PBS) 0.05% Tween 80 buffer (pH 7.4; *w/v*), resuspended in 200 µL sterile PBS and further sonicated for 180 sec using an ultrasonic bath to reduce bacterial aggregation. Bacterial suspension (5 μL) was spotted between a coverslip of 170 μm thickness and a freshly prepared 1.5% agarose-PBS pad of approximately 1-2 cm^2^. Bacteria were analysed by snapshot imaging at room temperature using a Leica SP5 laser scanning confocal microscope (Leica Biosystems). Image acquisition was performed with an HC PL APO CS2 63×/1.40 oil objective. Images of 1,024 x 1,024 pixels were acquired with argon 488 nm, and diode-pumped solid-state 561 nm lasers. Emitted signal was collected at λ_em_ 500-550 nm and λ_em_ 585-650 nm for mWasabi and dTomato channels respectively. One single Z-plane was acquired for each field, and a minimum of 5 fields per biological sample were imaged. Individual raw fluorescence images were exported as TIFF files and displayed with open-source software such as Fiji/imagej software (https://imagej.net/software/fiji/) [31]. Visualisation was performed at least on two-independent occasions.

## Results and Discussion

### Generation of a new subset of fluorescence-based CRISPRi vectors

In order to generate new fluorescent CRISPRi vectors derived from the original pIRL series, we selected fluorophores that have been previously reported for their brightness and stability in mycobacteria [25]. As such, we postulated that these new vectors should display strong fluorescent signals upon recombination at the mycobacteriophage L5 integration site, located at the tRNA^gly^ gene [11, 35]. We chose the well-established and widely used mWasabi (λ_Ex_/λ_Em_ 490/510 nm) and dTomato (λ_Ex_/λ_Em_ 550/580 nm) fluorophores previously described in the Ramakrishnan group [25].

Since the sgRNA targeting sequence cloning strategy in the pIRL series is based on the use of the BsmB1 enzyme and its corresponding restrictions sites [10, 11], the presence of the BsmB1 restriction site within the upstream promoter region of the genes in the pTEC15 and pTEC27 vectors led us to subclone the *mWasabi* and *dTomato* genes in an alternative vector. Thus, *mWasabi* and *dTomato* sequences were PCR amplified and further cloned at the SphI and HindIII restriction sites of the pUV15-pHGFP vector (Addgene plasmid #70045) [29] to replace the pH-GFP encoding gene (Fig.S1A). Using this strategy, two new episomal vectors carrying *mWasabi* or *dTomato* genes under the control of the strong and constitute *psmyc* promotor were generated and respectively named pJL29 and pJL30 (Fig.S1A).

Recombinant *M. smegmatis* strains harbouring pJL29 and pJL30 were generated alongside *M. smegmatis* pMV306-Hyg, pTEC15, pTEC27 controls strains (Fig.S1B & Fig.S1C). Fluorescence signals derived from these strains were qualitatively assessed on both agar media and in liquid culture (Fig.S1D & Fig.S1E). These new constructs displayed strong fluorescence signals with intensities very close to the original pTEC15 and pTEC27 vectors, suggesting that they can be used for the subsequent development of pIRL-derivative vectors (Fig.S1D & Fig.S1E).

To generate the fluorescent pIRL-derivatives, *mWasabi* and *dTomato* genes were PCR amplified with their upstream *psmyc* promotor from the pJL29 and pJL30 plasmids, and subsequently cloned as blunt-end fragments in HpaI-digested pIRL2 and pIRL117 (Addgene plasmids #163631 & #163635) [11]. Recombinant clones were selected and successful cloning of the fluorescent markers was checked by whole plasmid sequencing. Hence, the newly generated vectors derived from pIRL117 were named as pJL31 (mWasabi) and pJL32 (dTomato) and the ones derived from pIRL2 were named pJL33 (mWasabi) and pJL34 (dTomato), respectively (Fig.1A).

**Figure 1.**
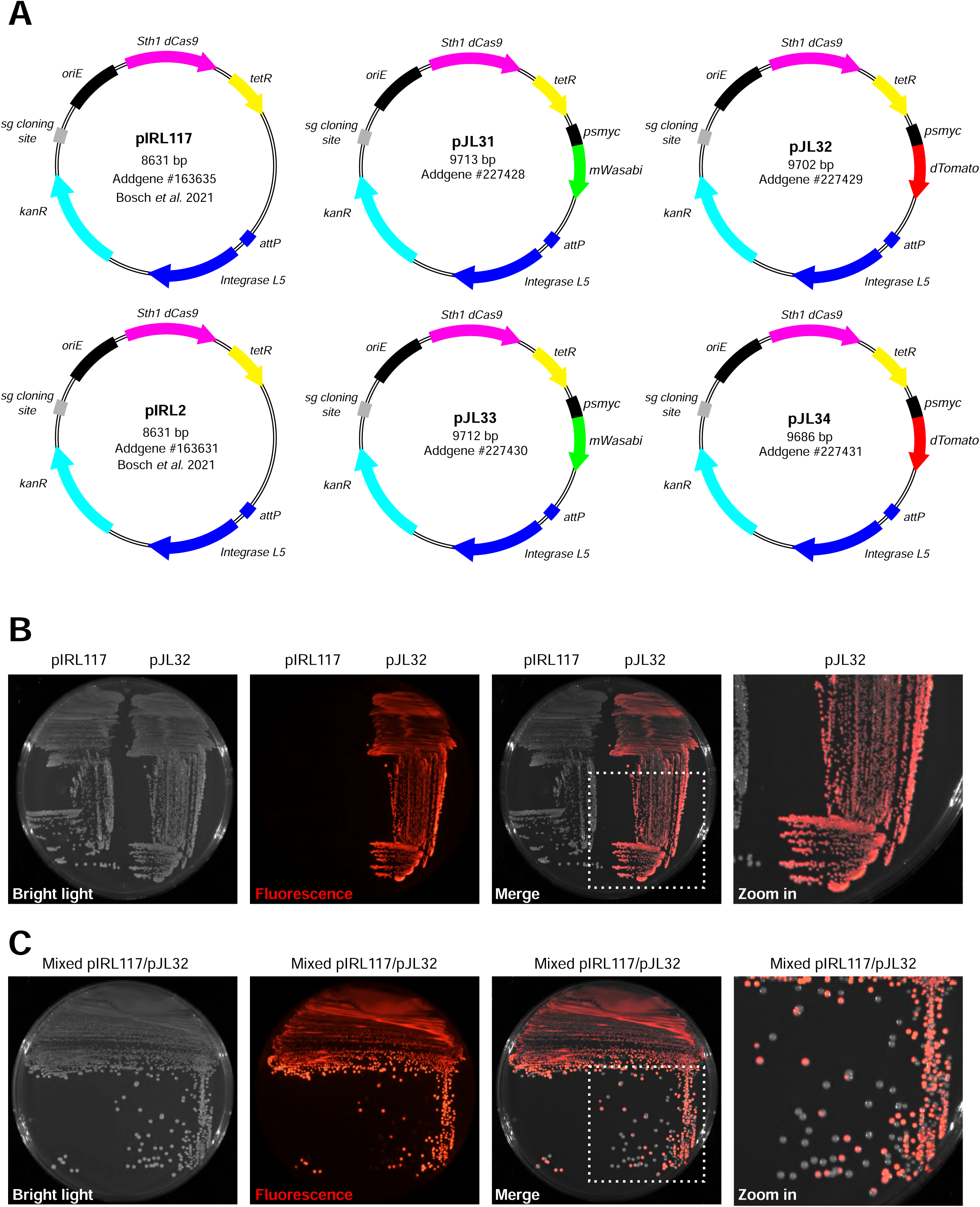
Generation of a new series of fluorescence-based CRISPRi vectors. **(A)** Schematic representation of the original pIRL2-pIRL117 CRISPRi integrative vectors and the newly generated pJL31, pJL32, pJL33 and pJL34 derivatives harbouring the *mWasabi* or *dTomato* coding sequence under the control of the strong constitutive *psmyc* promotor. **(B)** Detection and comparative analysis of *M. smegmatis* recombinant strains harbouring the original pIRL117 construct or the pJL32 vector when plated onto 7H10 agar plates. Bright light, red fluorescence, merge micrographs are displayed from left to right. The far-right micrograph corresponds to an additional Zoom caption of the Merge micrographs. **(C)** Mixed population (50%/50%) of *M. smegmatis* pIRL117 and pJL32 recombinant strains were plated onto 7H10 agar plates and separated using the quadrant streak plate technique. Bright light, red fluorescence, merge micrographs are displayed from left to right. The far-right micrograph corresponds to an additional Zoom caption of the Merge micrographs.

To further validate that such vectors can be used to easily identify and select recombinant mycobacterial clones that have integrated CRISPRi vectors at the *attB* chromosomal attachment site, we generated recombinant *M. smegmatis* strains harbouring the pIRL117 or the pJL32 vectors. Detection and comparison of fluorescent and non-fluorescent strains was performed following inoculation and streaking on agar plates (Fig.1B). As proof of concept, recombinant strains were spotted as macro-colonies (Fig.S2A) or cultured in broth (Fig.S2B) to show that a simple bench-top imaging system can easily detect fluorescent recombinants clones from their non-fluorescent counterparts. Final validation was performed by mixing equally pIRL117 and pJL32 recombinant strains and further isolating them by using the quadrant streak plate technique (Fig.2C). Results displayed Fig.2C demonstrate that integration of pJL32 enables a simple a rapid discrimination from other kanamycin resistant colonies, thus facilitating the selection and expansion process after transformation.

**Figure 2.**
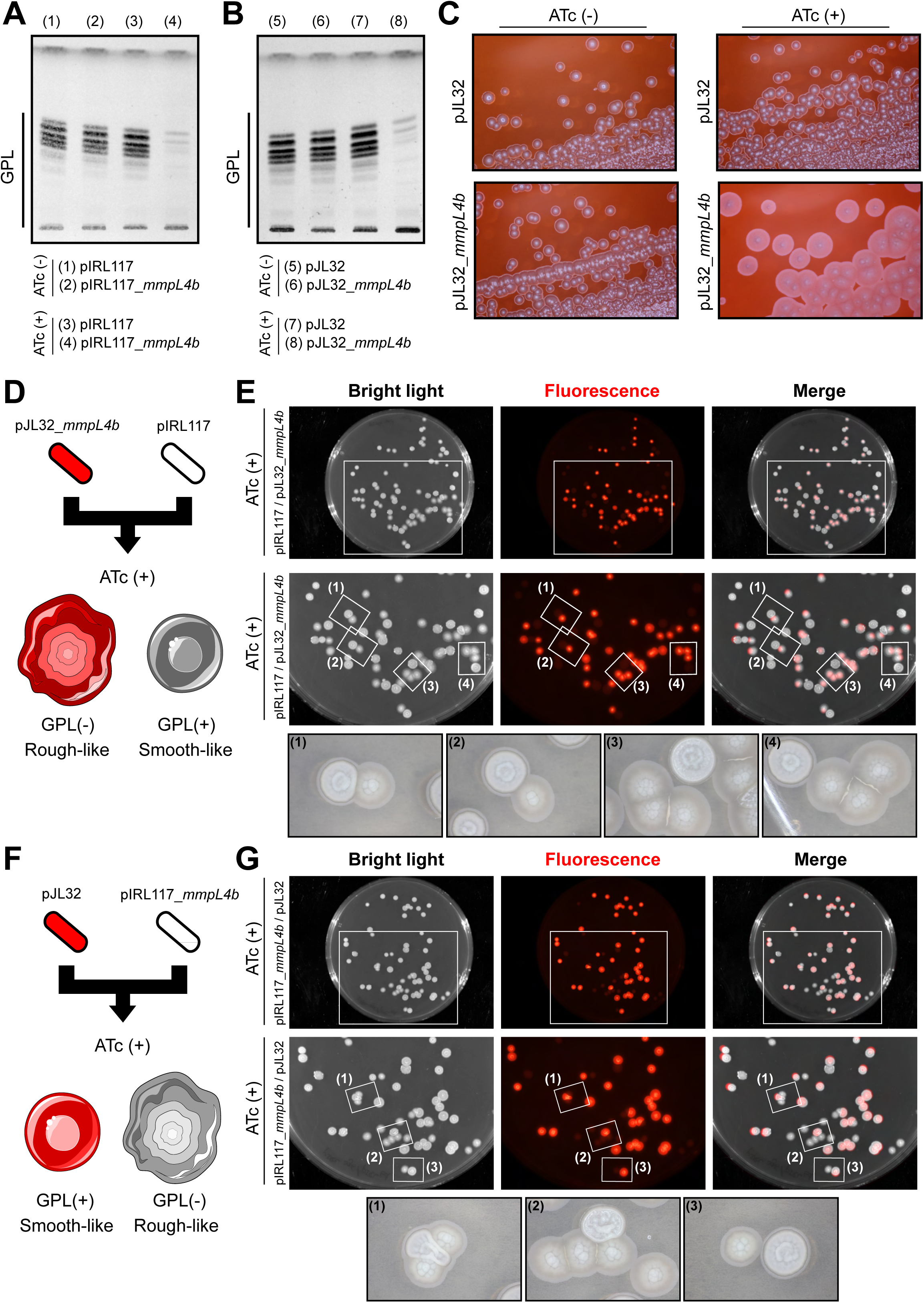
Functional validation of fluorescence-based CRISPRi vectors by targeting the non-essential mycobacterial GPL translocase MmpL4. **(A)** GPL analysis of *M. smegmatis* pIRL117 or pIRL117_*mmpL4b* in the presence or in the absence of ATc. Cells were grown in 7H9 broth +/− 100 ng/mL of ATc for 48 h. Lipid extraction from normalized dry pellets was analyzed by TLC. Lanes are numbered from (1) to (4) and the corresponding loaded samples are indicated at the bottom of the TLC plate. **(B)** GPL analysis of *M. smegmatis* pJL32 or pJL32_*mmpL4b* in the presence or in the absence of ATc. Cells were grown in 7H9 broth +/− 100 ng/mL of ATc for 48 h. Lipid extraction from normalized dry pellets was analyzed by TLC. The loaded samples are indicated at the bottom of the TLC plate (Lanes (5) to (8)). **(C)** Congo-red binding evaluation of *mmpL4b* CRISPRi silencing using the pJL32 and pJL32_*mmpL4b* vectors. Cells were grown in 7H10 solid medium containing 100 µg/mL of Congo-red dye +/− 100 ng/mL of ATc for 48 h before being imaged using a stereomicroscope coupled with a Canon Digital Camera. **(D)** Schematic representation of the mixing experiments with control pIRL117 and pJL32_*mmpL4b* vectors in the presence of ATc with their respective expected phenotypes. **(E)** Analysis of *M. smegmatis* pJL32_*mmpL4b* / pIRL117 morphologies and fluorescence profiles when mixed on solid medium. Approximately, 50 CFU of *M. smegmatis* pJL32_*mmpL4b* and 50 CFU of *M. smegmatis* pIRL117 were mixed before being plated on 7H10 solid medium containing 100 ng/mL of ATc. After 3-4 days at 37°C the plates were imaged using bench-top imager. Bright light, red fluorescence, merge micrographs are displayed from left to right. Zoom caption of each channel are displayed in the middle and selected area from (1) to (4) have been further imaged using a stereomicroscope coupled with a Canon Digital Camera. Their corresponding micrographs are showed at the bottom of the panel. **(F)** Schematic representation of the mixing experiments with control pJL32 and pIRL117_*mmpL4b* vectors in the presence of ATc with their respective expected phenotypes. **(G)** Analysis of *M. smegmatis* pIRL117_*mmpL4b* / pJL32 morphologies and fluorescence profiles when mixed on solid medium. Experimental setting was exactly the same as indicated in **(E)**.

### Functional validation of fluorescent-based CRISPRi vectors through targeted repression of the non-essential GPL translocase *mmpL4b*

To validate that the newly generated vectors have fully conserved their ability to achieve inducible targeted repression, we performed comparative analysis of pIRL117-mediated repression with pJL32-mediated repression. To do so, we first cloned a 21-bp sgRNA targeting sequence that targets the non-template strand of the *mmpL4b* gene in both pIRL117 and the newly generated pJL32 as previously described [9–11] (Table S1). The *mmpL4b* gene is involved in the proton-dependent translocation of glycopeptidolipids (GPL) from the cytosol to the cell-wall of non-tuberculous mycobacteria [36–38]. This gene was chosen because the presence of GPL in the mycobacterial envelope can be easily evaluated by TLC analysis or assessed through morphological analysis of the bacterial colonies on conventional agar medium [20, 37, 39].

Recombinant strains harbouring pIRL117, pIRL117_*mmpL4b*, pJL32 or pJL32_*mmpL4b* were grown in 7H9 broth in the presence or in the absence of 100 ng/mL of ATc for 48 h, and GPL levels were analysed by TLC analysis (Fig.2A & 2B). Comparison of *M. smegmatis* pIRL117 and pIRL117_*mmpL4b* GPL content in the absence of inducer did not show any differences, supporting the fact that no major leaky expression of sgRNA-dCas9 occurred (Fig.2A - lanes (1) & (2)). In addition, GPL profiles of the *M. smegmatis* pIRL117 control strain in the presence or absence of ATc were similar, suggesting that ATc treatment does not impact GPL production/translocation (Fig.2A - lanes (1) & (3)). Finally, analysis of *M. smegmatis* pIRL117_*mmpL4b* profiles demonstrated that upon ATc treatment GPL content from the extractible fraction was almost null, confirming a fully-functional repression system of the *mmpL4b* gene (Fig.2A - lanes (2) & (4)).

To validate our newly generated tools, the same experiments were carried out with the pJL32 or pJL32_*mmpL4b* recombinant strains. Results showed that the exact same profiles were obtained, where *mmpL4b* gene repression was ATc-dependent and only observed in the pJL32_*mmpL4b* strain but not the pJL32 control strain (Fig.2B).

Since GPL production is associated with important colony morphological changes, where GPL^(+)^ strains display smooth-like phenotypes (S morphotype) while GPL^(-)^ strain often have rough-like phenotype (R morphotype) [38, 40, 41]. We wanted to confirm our results obtained previously in liquid cultures and demonstrate that our repression system allows to distinguish distinct morphotypes on solid medium (Fig.2C). To do so, we plated both *M. smegmatis* pJL32 and pJL32_*mmpL4b* on 7H10 solid media containing the cell-wall binding Congo-Red dye [32, 33]. Morphological features of the colony in the presence or absence of 100 ng/mL ATc were assessed after 48 hours of growth using a stereomicroscope. Results demonstrated that *M. smegmatis* pJL32 colonies remained small, smooth and shiny regardless ATc levels. On the other hand, *M. smegmatis* pJL32_*mmpL4b* colonies displayed a bigger, drier and less regular morphotype in the presence of ATc while the control experiment devoid of ATc resulted in phenotypes similar to the empty pJL32.

Finally, we set-up another experimental setting by performing mixing experiments of *M. smegmatis* strains harbouring the empty pIRL117 and the pJL32_*mmpL4b* (Fig.2D-E) or the other way around by mixing strains harbouring the empty pJL32 and the pIRL117_*mmpL4b* (Fig.2F-G). Following this experimental scheme, plating pIRL117 and pJL32_*mmpL4b* mixed populations on solid agar media containing 100 ng/mL of ATc should lead to the formation of a rough-like morphotype only for the fluorescent colonies (Fig.2D). Results displayed in Fig.2E confirmed that only the red-fluorescent colonies were the one displaying rough-like phenotypes while the control non-fluorescent bacteria conserved a shiner, more regular smooth-like phenotype as expected (Fig.2E). Complementary experiments with the opposite combination were also performed, where the non-fluorescent pIRL117_*mmpL4b* and the pJL32, without sgRNA targeting sequence were used (Fig.2F). Results presented in the Fig.2G fully confirmed our findings, with rough-like morphotypes only found this time in the non-fluorescent *M. smegmatis*.

Altogether, the findings obtained from this new subset of experiments confirmed the results previously observed in broth and Congo-Red containing plates, further demonstrating that our system is fully functional in multiple experimental settings.

### Functional validation of fluorescent-based CRISPRi vectors through targeted repression of the essential trehalose monomycolate translocase *mmpL3* and the β-subunit of the RNA-polymerase *rpoB*

Since CRISPRi is now widely used to investigate the function of essential genes, we decided to validate our fluorescent constructs by targeting essential genes. For that, the trehalose monomycolate translocase *mmpL3* and the β-subunit of the RNA-polymerase *rpoB* were selected as candidate genes. Both sgRNA targeting sequences were cloned in the pIRL117 and the pJL32 vectors. Single colonies of all the generated strains carrying the pIRL117 or pJL32-derivatives were used to inoculate 7H9 broth, and fluorescence levels were checked using our bench top imaging system (Fig.3A).

**Figure 3.**
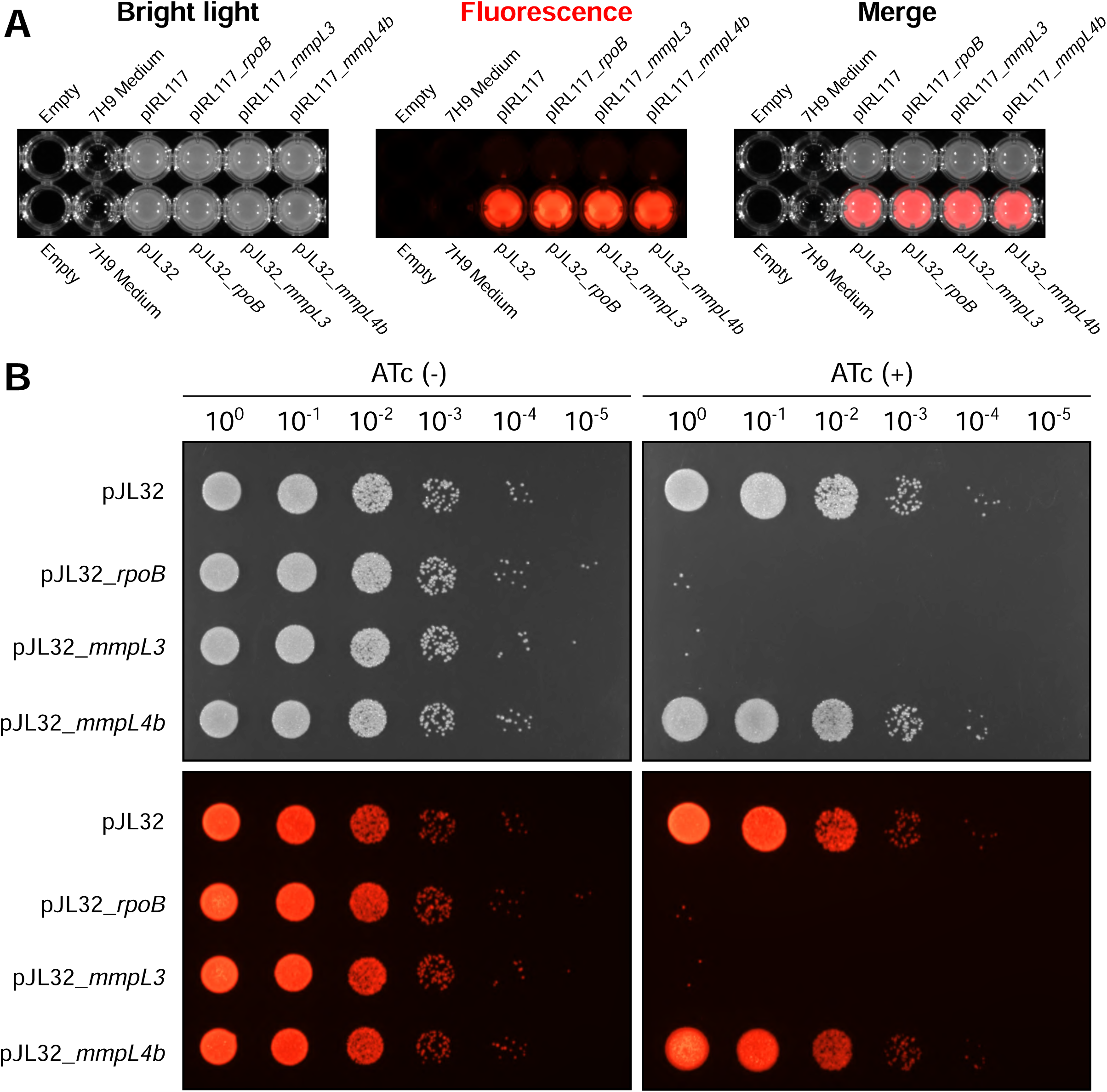
Functional validation of fluorescence-based CRISPRi vectors by targeting essential mycobacterial genes. **(A)** Fluorescence display of *M. smegmatis* pIRL117 and pJL32 recombinant strains when cultured in 7H9 liquid broth. Strains harbouring the pIRL117 and pJL32 empty vectors or with the *rpoB*, *mmpL3*, and *mmpL4b* targeting sequences were imaged using the fluorescence imager. Bright light, red fluorescence, merge micrographs are displayed from left to right. **(B)** Functional validation of pJL32 CRISPRi system by targeting *rpoB*, *mmpL3*, and *mmpL4b*. Serial dilution of *M. smegmatis* recombinants strains was spotted onto 7H10 agar media in the absence ATc (left panel) or in the presence ATc (right panel) of 100 ng/mL of anhydrotetracycline. Bright light (top) and their corresponding red fluorescent profiles (bottom) are displayed.

Then, serial dilution of each pJL32 recombinant strains were spotted onto 7H10 agar plates in the presence or absence of 100 ng/mL of ATc (Fig.3B). Plating on standard plates without ATc did not show any major modifications in between the genetic background (Fig.3B; left panel). However, serial spotting on ATc-containing plates resulted in an approximate 4-log^10^ decrease in bacterial growth suggesting that both vectors were fully functional (Fig.3B; right panel). No major defects were observed for the strains carrying the empty pJL32 and the pJL32_*mmpL4b,* which have no target or targets a non-essential gene respectively (Fig.3B; right panel). This experiment was then repeated with both pJL32 and non-fluorescent pIRL117-derivatives as comparative control and similar findings were obtained (Fig.S3), further confirming that the pJL series is perfectly suitable for performing CRISPRi-mediated silencing.

### One-step co-transformation of fluorescent-based CRISPRi vectors with alternative fluorescence reporters to enable genetic studies coupled with single-cell dual imaging

Since genetic modification through targeted mutagenesis and the subsequent transformation of fluorescent-vectors of interest in a mutant genetic background could be time-consuming, we thought about testing the potential of our new constructs to be simultaneously co-electroporated with additional vectors. Indeed, pJL29 and pJL30 being episomal vectors and conferring hygromycin resistance, they could be perfectly compatible with the use of the pJL31, pJL32, pJL33 or pJL34 which are integrative and conferring kanamycin resistance. As such, a fluorescent genetic tool and an alternative fluorescent reporter could be potentially co-electroporated in mycobacteria in a single-step, therefore constituting a real advantage for subsequent studies.

This strategy was first tested on *M. smegmatis*, in which competent cells were transformed with the pJL29, the pJL32 or both of them simultaneously, and the recombinant transformants selected on agar plates containing 50 mg/L of kanamycin, 50 mg/L of hygromycin or 50 mg/L of both respectively (Fig.4A). Micrographs displayed in Fig.4A demonstrate that such strategy is perfectly suitable to obtain co-transformants in a simple one-step procedure. Fluorescence profiles were also assessed on agar as microcolonies or in liquid media (Fig.4B). We further investigated whether this approach could be used for strict pathogen such as *M. tuberculosis*. Similar experiments were performed using the pJL29/pJL34 vectors and co-transformants were also obtained post-electroporation for *M. tuberculosis* (Fig.S4). Therefore, we propose that such strategy, which is extremely easy to implement, could be complementary used alongside previously published methodologies to facilitate bacterial genetics and mycobacterial research [19, 42].

**Figure 4.**
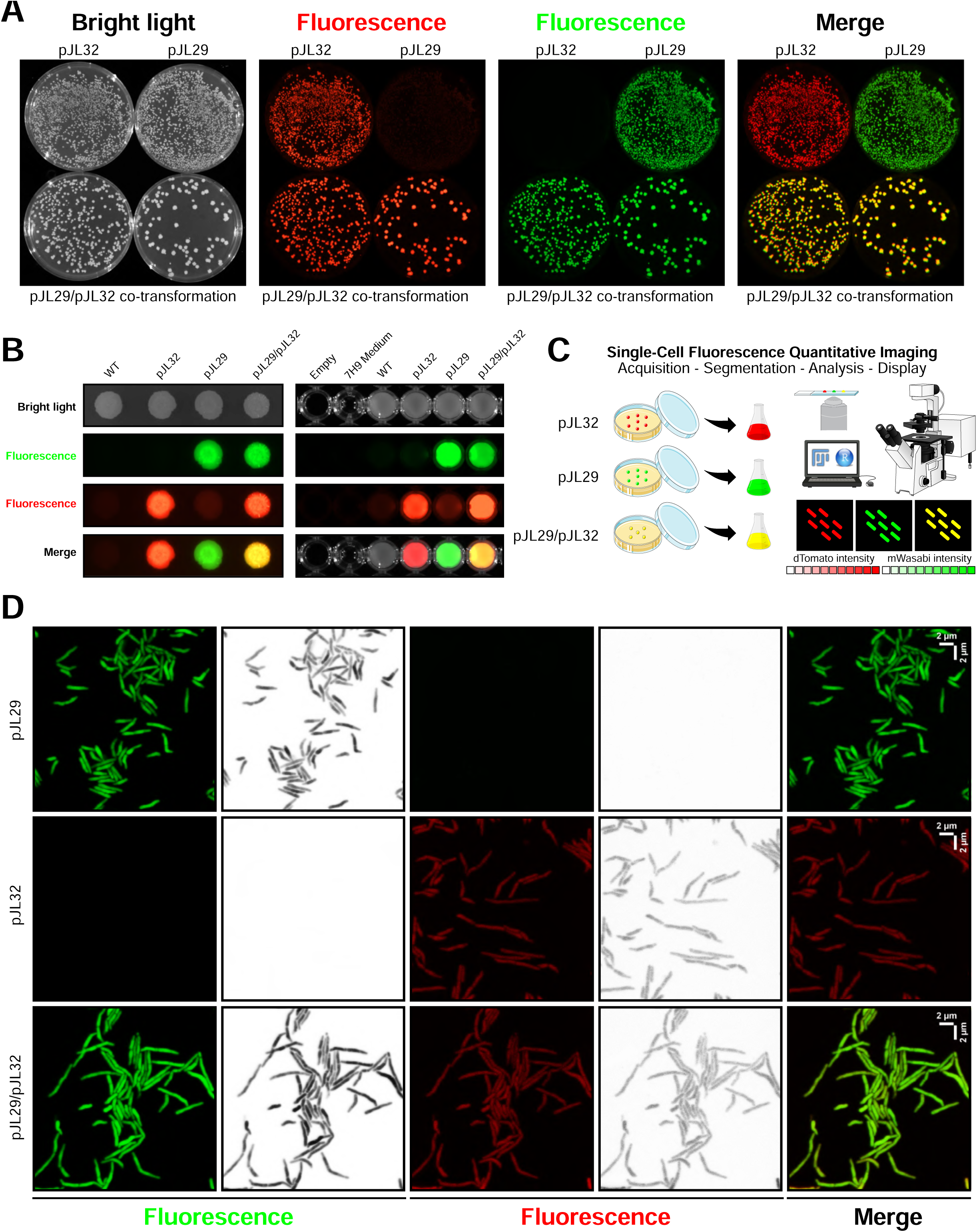
Fluorescent-based CRISPRi vectors can be associated with alternative reporters through a simple one-step co-transformation/selection procedure and imaged live at the single-cell level. **(A)** Fluorescence display of *M. smegmatis* pJL29, pJL32 or pJL29/32 recombinant strains when selected post-electroporation onto 7H10 agar plates. Bright light, red fluorescence, green fluorescence and merge micrographs are displayed from left to right. **(B)** Fluorescence display of *M. smegmatis* pJL29, pJL32 or pJL29/pJL32 recombinant strains when cultured as macro-colonies on 7H10 solid media (left) or in 7H9 liquid broth (right). Bright light, green fluorescence, red fluorescence and merge micrographs are displayed from top to bottom. **(C)** Schematic representation of the single-cell fluorescence microscopy assays that can be used when co-transforming reporters. **(D)** High-resolution fluorescence imaging of *M. smegmatis* bacterial cells harbouring pJL29, pJL32 or both pJL29/pJL32. Cells were imaged in both green (left panel; in green and in black and white) and red fluorescent channels (middle panel; in red and in black and white). Merged micrographs are displayed on the right. Scale bars represent 2 µm in X and Y.

Finally, we postulated that such co-transformation strategy could be very useful to investigate gene functions by targeted repression using dual single-cell imaging (Fig.5C). To test this, *M. smegmatis* transformants or co-transformants were imaged live using fluorescence microscopy (Fig.5D). Results showed that recombinant strains harbouring the pJL29 episomal vectors were displaying strong and easily detectable fluorescence profiles. On the other hand, recombinant strains carrying the pJL32 integrative vector, was as expected, less bright but still showing a good signal-to-noise ratio, and giving the possibility to distinguish individual bacterial cell. Co-transformants, with the expression of the two-fluorophores could be imaged very easily and showed signals similar to the ones obtained from single transformants (Fig.5D). This new observation really opens new possibilities regarding genetic modifications and live single-cell imaging. Indeed, one can easily imagine using the CRISPRi pJL series to repress gene expression and rely on its derived fluorescence as control and/or segmentation channel. Then, an alternative fluorophore such as a biosensor or any translational/transcriptional fusion could be used to further investigate new molecular function(s)/mechanism(s) directly related to the targeted candidate gene, thus enabling new cutting-edge investigations with the ultimate goal of making ground-breaking discoveries.

### Concluding remarks

In this letter, we have reported the development and functional validation of fluorescent-based CRISPRi vectors. The latter are available in two colours (mWasabi/green or dTomato/red) and have been developed for both fast and slow growing species by modifying the two original vectors pIRL117 and pIRL2. Therefore, these tools should simplify the selection process of CRISPRi recombinants and facilitate any subsequent investigations that might require fluorescence-based quantification. In addition, we have reported that these tools are compatible with episomal vectors carrying alternative fluorophores or potentially any gene of interest, and that such combination could be easily obtained through a single round of co-electroporation, highlighting the strong potential of this novel approach. Overall, we believe that making these tools fully available for our community through non-profit depositories should help designing new experimental approaches that have the potential to foster new discoveries in mycobacterial research.

## Abbreviations

(CRISPRi): CRISPR interference
(CFU): colony forming units
(OD_600nm_): optical density 600 nm
(ATc): anhydrotetracycline
(GPL): glycopeptidolipids
(PBS): phosphate buffer saline
(sgRNA): single-guide ribonucleic acid

## Acknowledgements

We would like to acknowledge all members of the Lipolysis and Bacterial Pathogenicity group and the LISM unit for continuous support and insightful discussions. PS would like to specially thank Max Gutierrez for his continuous support, help and mentorship over the years.

This work was supported by the Centre National de la Recherche Scientifique (CNRS) and Aix-Marseille Université (AMU). PS received financial support from the CNRS Biologie, the Agence Nationale de Recherches sur le Sida et les Hépatites virales (ANRS) (project n°ANRS0358) and the French government under the France 2030 investment plan, as part of the Initiative d’Excellence d’Aix-Marseille Université - A*MIDEX and is part of the Institute of Microbiology, Bioenergies and Biotechnology - IM2B (AMX-19-IET-006). JL PhD fellowship was funded by the Ministère de l’Enseignement Supérieur et de la Recherche. WA postdoctoral fellowship was funded by the foundation IHU Méditerranée Infection.

PS has also received a FEBS Excellence Award 2023 to support this work and would like to personally thank the FEBS Fellowships Office, FEBS Letters and FEBS Open Bio Editorial Offices for their continuous support.

The funders did not play a role in the study design, data collection and analysis, decision to publish, or preparation of the manuscript.

## Authors contributions

PS proposed, conceived and led the study. PS secured funding. SC co-advised the PhD work of JL with PS. SC and CC co-advised the postdoctoral work of WA with PS. JL, VP, WA and PS performed the experimental work. JL, VP and PS edited figures. All authors provided intellectual input by organising, analysing, and/or discussing data. PS wrote the manuscript. All authors read the manuscript and provided critical feedback before its submission.

## Data availability statement

Plasmids vectors from the pJL series generated in this study are available at https://www.addgene.org/.

Any additional data that support the findings of this study are available upon reasonable request from the corresponding author at.

## Competing interests

The authors declare no competing interests.

## Supporting information

**Table S1.**
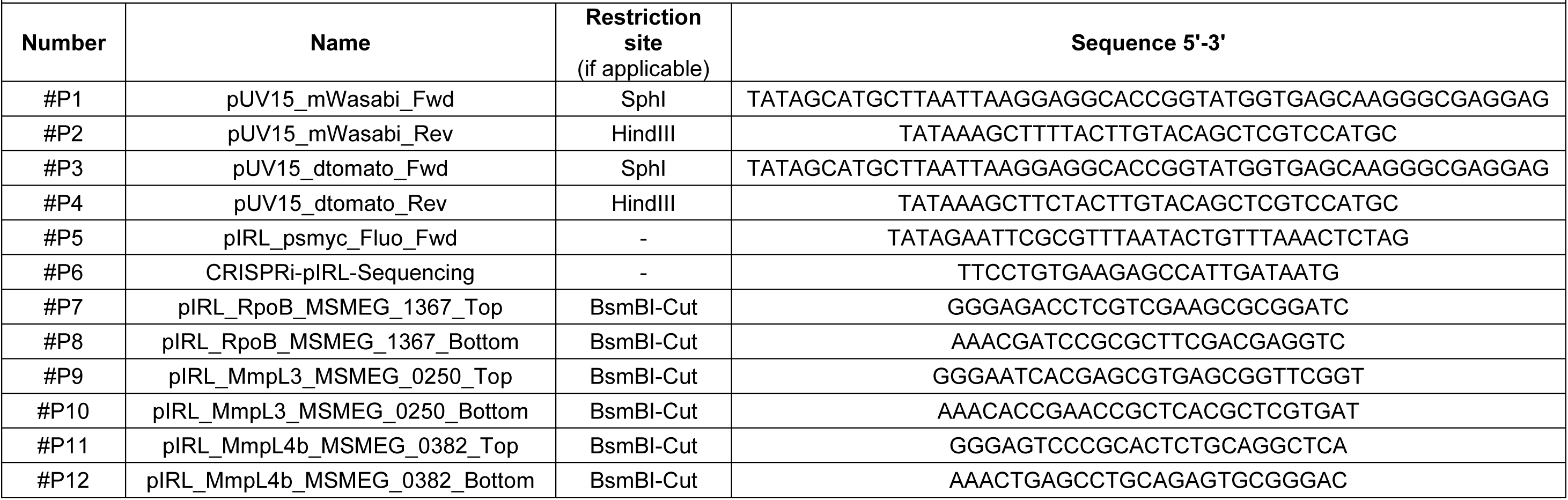
List of the primers used in this study.

**Table S2.** List of the plasmids used in this study.

| **Table S2. Plasmids used in this study** |  |  |  |
| --- | --- | --- | --- |
| **Name** | **Alternative Name** | **Short Description** | **Reference and/or Source** |
| pTEC15 | - | Hygromycin resistant episomal vector for the constitutive expression of mWasabi under the control of the *MSP12* promotor (pMSP12::mWasabi) | Takaki *et al.* 2013 - Addgene Plasmid #30174 |
| pTEC27 | - | Hygromycin resistant episomal vector for the constitutive expression of dTomato under the control of the *MSP12* promotor (pMSP12::dTomato) | Takaki *et al.* 2013 - Addgene Plasmid #30182 |
| pUV15-pHGFP | - | Hygromycin resistant episomal vector for the constitutive expression of pHGFP under the control of the *psmyc* promotor (psmyc::pHGFP) | Vandal *et al.* 2008 - Addgene Plasmid #70045 |
| pJL29 | pUV15_mWasabi | Hygromycin resistant episomal vector for the constitutive expression of mWasabi under the control of the *psmyc* promotor (psmyc::mWasabi) | This study - Addgene Plasmid #227426 |
| pJL30 | pUV15_dTomato | Hygromycin resistant episomal vector for the constitutive expression of dTomato under the control of the *psmyc* promotor (psmyc::dTomato) | This study - Addgene Plasmid #227427 |
| pIRL2 | - | Kanamycin resistant L5-integrative vector for CRISPR interference in *M.tuberculosis* | Bosch *et al.* 2020 - Addgene Plasmid #163631 |
| pIRL117 | - | Kanamycin resistant L5-integrative vector for CRISPR interference in *M.smegmatis* | Bosch *et al.* 2020 - Addgene Plasmid #163635 |
| pJL31 | pIRL117_*psmyc::mWasabi* | Kanamycin resistant L5-integrative vector for CRISPR interference in *M.smegmatis* with constitutive expression of mWasabi under the control of the *psmyc* promotor (psmyc::mWasabi) | This study - Addgene Plasmid #227428 |
| pJL32 | pIRL117_*psmyc::dTomato* | Kanamycin resistant L5-integrative vector for CRISPR interference in *M.smegmatis* with the constitutive expression of dTomato under the control of the *psmyc* promotor (psmyc::dTomato) | This study - Addgene Plasmid #227429 |
| pJL33 | pIRL2_*psmyc::mWasabi* | Kanamycin resistant L5-integrative vector for CRISPR interference in *M.tuberculosis* with the constitutive expression of mWasabi under the control of the *psmyc* promotor (psmyc::mWasabi) | This study - Addgene Plasmid #227430 |
| pJL34 | pIRL2_psmyc::dTomato | Kanamycin resistant L5-integrative vector for CRISPR interference in *M.tuberculosis* with the constitutive expression of dTomato under the control of the psmyc promotor (psmyc::dTomato) | This study - Addgene Plasmid #227431 |
| pJL1 | pIRL117_rpoB | Kanamycin resistant L5-integrative vector for CRISPR interference of *rpoB* in *M.smegmatis* | This study |
| pJL2 | pIRL117_mmpL3 | Kanamycin resistant L5-integrative vector for CRISPR interference of *mmpL3* in *M.smegmatis* | This study |
| pJL3 | pIRL117_mmpL4b | Kanamycin resistant L5-integrative vector for CRISPR interference of *mmpL4b* in *M.smegmatis* | This study |
| pJL35 | pJL32_rpoB | Kanamycin resistant L5-integrative vector for CRISPR interference of *rpoB* in *M.smegmatis* with the constitutive expression of dTomato under the control of the psmyc promotor (psmyc::dTomato) | This study |
| pJL36 | pJL32_mmpL3 | Kanamycin resistant L5-integrative vector for CRISPR interference of *mmpL3* in *M.smegmatis* with the constitutive expression of dTomato under the control of the psmyc promotor (psmyc::dTomato) | This study |
| pJL37 | pJL32_mmpL4b | Kanamycin resistant L5-integrative vector for CRISPR interference of *mmpL4b* in *M.smegmatis* with the constitutive expression of dTomato under the control of the psmyc promotor (psmyc::dTomato) | This study |

**Figure S1.**
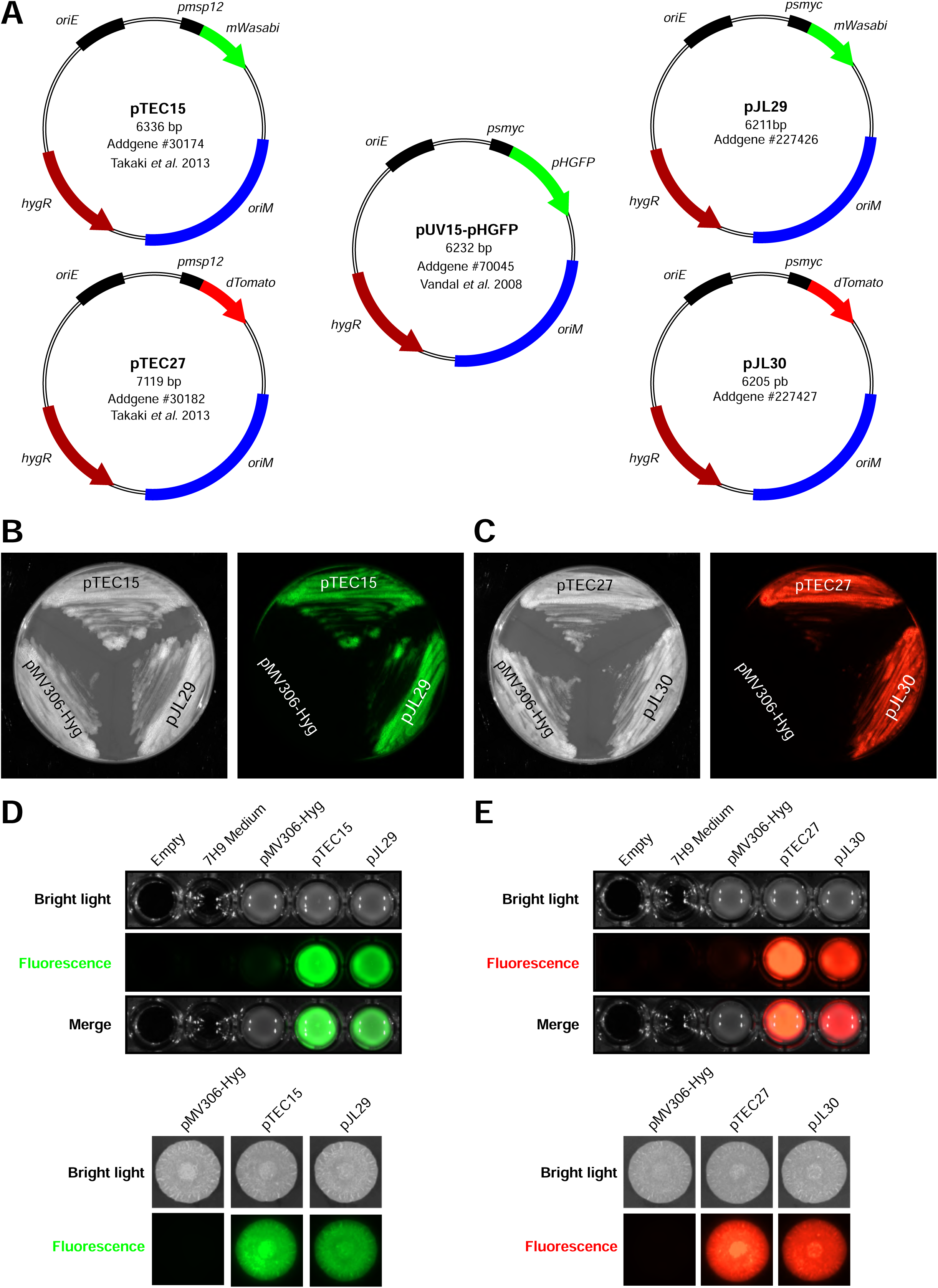
Generation and functional validation of pJL29 and pJL30 fluorescent vectors. **(A)** Schematic representation of the original pTEC15, pTEC27 and pUV15-pHGFP vectors and the newly generated pJL29 and pJL30 vectors harbouring the *mWasabi* or *dTomato* coding sequence under the control of the strong constitutive *psmyc* promotor. **(B-C)** Detection and comparative analysis of *M. smegmatis* recombinant strains harbouring the original pTEC15, pTEC27 construct or the pJL29 or pJL30 vectors when plated onto 7H10 agar plates. The pMV306-Hyg was used as fluorescent-negative control. Bright light, green fluorescence, red fluorescence, merge micrographs are displayed. **(D-E)** Analysis of pJL-mediated fluorescence in comparison to its parental pTEC vectors on agar and in liquid medium. Fluorescence display of *M. smegmatis* recombinant strains when spotted as macro-colonies on 7H10 agar plates (bottom panel) or when cultured in 7H9 broth media (top panel).

**Figure S2.**
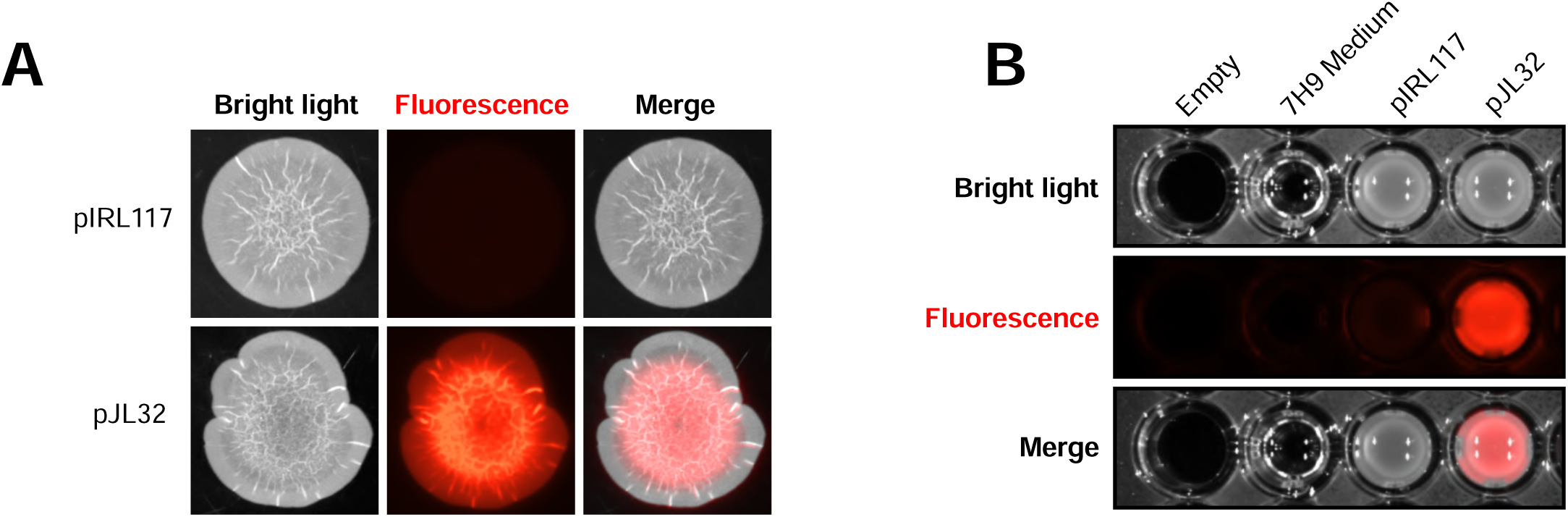
Analysis of pJL32-mediated fluorescence in comparison to its parental pIRL117 vector on agar and in liquid medium. **(A)** Fluorescence display of *M. smegmatis* pIRL117 and pJL32 recombinant strains when spotted as macro-colonies on 7H10 agar plates. **(B)** Fluorescence display of *M. smegmatis* pIRL117 and pJL32 recombinant strains when cultured in 7H9 broth media.

**Figure S3.**
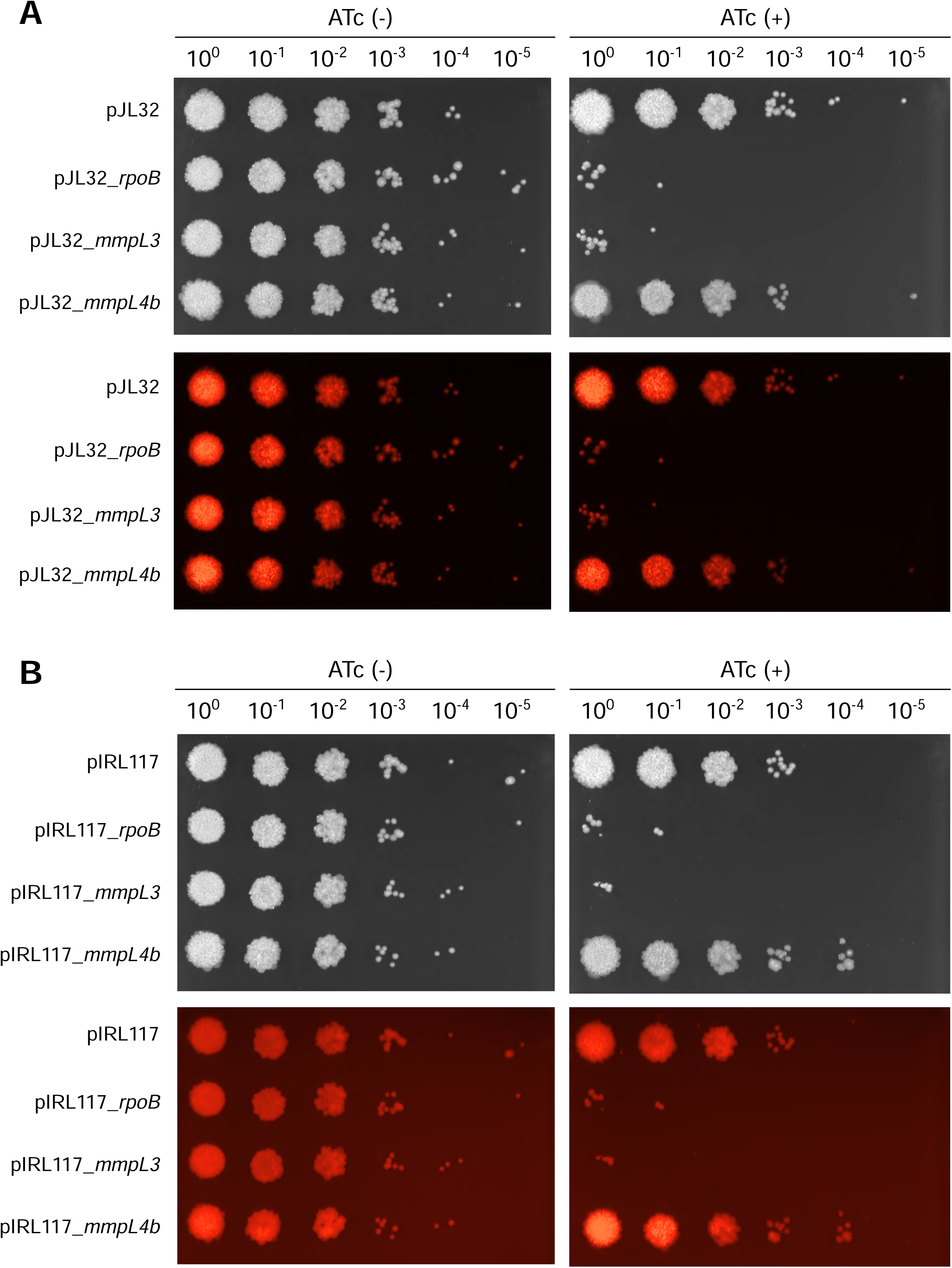
Comparison of pIRL117- and pJL32-mediated targeting of essential genes in *M. smegmatis*. **(A)** Functional validation of pJL32 CRISPRi system by targeting *rpoB*, *mmpL3*, and *mmpL4b*. Serial dilution of *M. smegmatis* recombinants strains was spotted onto 7H10 agar media in the absence ATc (Left panel) or in the presence ATc (Right panel) of 100 ng/mL of anhydrotetracycline. Bright light (top) and their corresponding red fluorescent profiles (bottom) are displayed. **(B)** Functional validation and comparison of pIRL117 CRISPRi system by targeting *rpoB*, *mmpL3*, and *mmpL4b*. Serial dilution of *M. smegmatis* recombinant strains was spotted onto 7H10 agar media in the absence ATc (Left panel) or in the presence ATc (Right panel) of 100 ng/mL of anhydrotetracycline. Bright light (top) and their corresponding red fluorescent profiles (bottom) are displayed. Non-fluorescent recombinant strains display a very low fluorescence signal which is as comparable as the background intensity from the medium.

**Figure S4.**
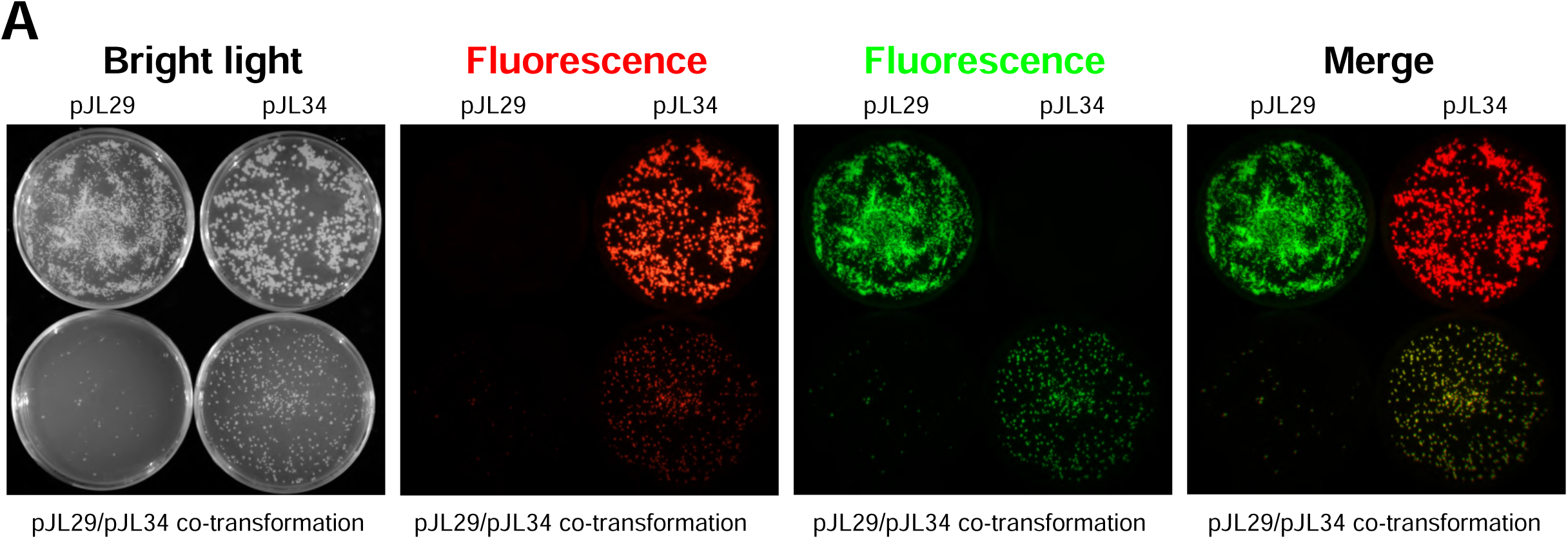
Illustration of the one-step co-transformation experiment performed in *M. tuberculosis*. **(A)** Fluorescence display of *M. tuberculosis* pJL29, pJL34 or pJL29/34 recombinant strains when selected post-electroporation onto 7H10 agar plates. Bright light, red fluorescence, green fluorescence and merge micrographs are displayed from left to right.

## Notes

### Competing Interest Statement

The authors have declared no competing interest.

## References

1. Sun, B., Yang, J., Yang, S., Ye, R. D., Chen, D. & Jiang, Y. (2018) A CRISPR-Cpf1-Assisted Non-Homologous End Joining Genome Editing System of Mycobacterium smegmatis, Biotechnol J. 13, e1700588.

2. Meijers, A. S., Troost, R., Ummels, R., Maaskant, J., Speer, A., Nejentsev, S., Bitter, W. & Kuijl, C. P. (2020) Efficient genome editing in pathogenic mycobacteria using Streptococcus thermophilus CRISPR1-Cas9, Tuberculosis (Edinb). 124, 101983.

3. Liu, K., Gao, Y., Li, Z. H., Liu, M., Wang, F. Q. & Wei, D. Z. (2022) CRISPR-Cas12a assisted precise genome editing of Mycolicibacterium neoaurum, N Biotechnol. 66, 61–69.

4. Neo, D. M., Clatworthy, A. E. & Hung, D. T. (2024) A dual-plasmid CRISPR/Cas9-based method for rapid and efficient genetic disruption in Mycobacterium abscessus, J Bacteriol. 206, e0033523.

5. Akter, S., Kamal, E., Schwarz, C. & Lewin, A. (2024) Gene knock-out in Mycobacterium abscessus using Streptococcus thermophilus CRISPR/Cas, J Microbiol Methods. 220, 106924.

6. Yan, M. Y., Li, S. S., Ding, X. Y., Guo, X. P., Jin, Q. & Sun, Y. C. (2020) A CRISPR-Assisted Nonhomologous End-Joining Strategy for Efficient Genome Editing in Mycobacterium tuberculosis, mBio. 11.

7. Choudhary, E., Thakur, P., Pareek, M. & Agarwal, N. (2015) Gene silencing by CRISPR interference in mycobacteria, Nat Commun. 6, 6267.

8. Singh, A. K., Carette, X., Potluri, L. P., Sharp, J. D., Xu, R., Prisic, S. & Husson, R. N. (2016) Investigating essential gene function in Mycobacterium tuberculosis using an efficient CRISPR interference system, Nucleic Acids Res. 44, e143.

9. Rock, J. M., Hopkins, F. F., Chavez, A., Diallo, M., Chase, M. R., Gerrick, E. R., Pritchard, J. R., Church, G. M., Rubin, E. J., Sassetti, C. M., Schnappinger, D. & Fortune, S. M. (2017) Programmable transcriptional repression in mycobacteria using an orthogonal CRISPR interference platform, Nat Microbiol. 2, 16274.

10. Wong, A. I. & Rock, J. M. (2021) CRISPR Interference (CRISPRi) for Targeted Gene Silencing in Mycobacteria, Methods Mol Biol. 2314, 343–364.

11. Bosch, B., DeJesus, M. A., Poulton, N. C., Zhang, W., Engelhart, C. A., Zaveri, A., Lavalette, S., Ruecker, N., Trujillo, C., Wallach, J. B., Li, S., Ehrt, S., Chait, B. T., Schnappinger, D. & Rock, J. M. (2021) Genome-wide gene expression tuning reveals diverse vulnerabilities of M. tuberculosis, Cell. 184, 4579–4592 e24.

12. Li, S., Poulton, N. C., Chang, J. S., Azadian, Z. A., DeJesus, M. A., Ruecker, N., Zimmerman, M. D., Eckartt, K. A., Bosch, B., Engelhart, C. A., Sullivan, D. F., Gengenbacher, M., Dartois, V. A., Schnappinger, D. & Rock, J. M. (2022) CRISPRi chemical genetics and comparative genomics identify genes mediating drug potency in Mycobacterium tuberculosis, Nat Microbiol. 7, 766–779.

13. Boeck, L., Burbaud, S., Skwark, M., Pearson, W. H., Sangen, J., Wuest, A. W., Marshall, E. K. P., Weimann, A., Everall, I., Bryant, J. M., Malhotra, S., Bannerman, B. P., Kierdorf, K., Blundell, T. L., Dionne, M. S., Parkhill, J. & Andres Floto, R. (2022) Mycobacterium abscessus pathogenesis identified by phenogenomic analyses, Nat Microbiol. 7, 1431–1441.

14. Pelicic, V., Reyrat, J. M. & Gicquel, B. (1996) Positive selection of allelic exchange mutants in Mycobacterium bovis BCG, FEMS Microbiol Lett. 144, 161–6.

15. Pelicic, V., Jackson, M., Reyrat, J. M., Jacobs, W. R., Jr., Gicquel, B. & Guilhot, C. (1997) Efficient allelic exchange and transposon mutagenesis in Mycobacterium tuberculosis, Proc Natl Acad Sci U S A. 94, 10955–60.

16. Pavelka, M. S., Jr. & Jacobs, W. R., Jr. (1999) Comparison of the construction of unmarked deletion mutations in Mycobacterium smegmatis, Mycobacterium bovis bacillus Calmette-Guerin, and Mycobacterium tuberculosis H37Rv by allelic exchange, J Bacteriol. 181, 4780–9.

17. Parish, T. & Stoker, N. G. (2000) Use of a flexible cassette method to generate a double unmarked Mycobacterium tuberculosis tlyA plcABC mutant by gene replacement, Microbiology (Reading). 146 (Pt 8), 1969–1975.

18. Gregoire, S. A., Byam, J. & Pavelka, M. S. (2017) galK-based suicide vector mediated allelic exchange in Mycobacterium abscessus, Microbiology (Reading). 163, 1399–1408.

19. Murphy, K. C., Nelson, S. J., Nambi, S., Papavinasasundaram, K., Baer, C. E. & Sassetti, C. M. (2018) ORBIT: a New Paradigm for Genetic Engineering of Mycobacterial Chromosomes, mBio. 9.

20. Viljoen, A., Gutierrez, A. V., Dupont, C., Ghigo, E. & Kremer, L. (2018) A Simple and Rapid Gene Disruption Strategy in Mycobacterium abscessus: On the Design and Application of Glycopeptidolipid Mutants, Front Cell Infect Microbiol. 8, 69.

21. Richard, M., Gutierrez, A. V., Viljoen, A., Rodriguez-Rincon, D., Roquet-Baneres, F., Blaise, M., Everall, I., Parkhill, J., Floto, R. A. & Kremer, L. (2019) Mutations in the MAB_2299c TetR Regulator Confer Cross-Resistance to Clofazimine and Bedaquiline in Mycobacterium abscessus, Antimicrob Agents Chemother. 63.

22. Wakamoto, Y., Dhar, N., Chait, R., Schneider, K., Signorino-Gelo, F., Leibler, S. & McKinney, J. D. (2013) Dynamic persistence of antibiotic-stressed mycobacteria, Science. 339, 91–5.

23. Sommer, R. & Cole, S. T. (2019) Monitoring Tuberculosis Drug Activity in Live Animals by Near-Infrared Fluorescence Imaging, Antimicrob Agents Chemother. 63.

24. Fearns, A., Greenwood, D. J., Rodgers, A., Jiang, H. & Gutierrez, M. G. (2020) Correlative light electron ion microscopy reveals in vivo localisation of bedaquiline in Mycobacterium tuberculosis-infected lungs, PLoS Biol. 18, e3000879.

25. Takaki, K., Davis, J. M., Winglee, K. & Ramakrishnan, L. (2013) Evaluation of the pathogenesis and treatment of Mycobacterium marinum infection in zebrafish, Nat Protoc. 8, 1114–24.

26. Aylan, B., Botella, L., Gutierrez, M. G. & Santucci, P. (2023) High content quantitative imaging of Mycobacterium tuberculosis responses to acidic microenvironments within human macrophages, FEBS Open Bio. 13, 1204–1217.

27. Zhu, J., Wolf, I. D., Dulberger, C. L., Won, H. I., Kester, J. C., Judd, J. A., Wirth, S. E., Clark, R. R., Li, Y., Luo, Y., Gray, T. A., Wade, J. T., Derbyshire, K. M., Fortune, S. M. & Rubin, E. J. (2021) Spatiotemporal localization of proteins in mycobacteria, Cell Rep. 37, 110154.

28. de Wet, T. J., Winkler, K. R., Mhlanga, M., Mizrahi, V. & Warner, D. F. (2020) Arrayed CRISPRi and quantitative imaging describe the morphotypic landscape of essential mycobacterial genes, Elife. 9.

29. Vandal, O. H., Pierini, L. M., Schnappinger, D., Nathan, C. F. & Ehrt, S. (2008) A membrane protein preserves intrabacterial pH in intraphagosomal Mycobacterium tuberculosis, Nat Med. 14, 849–54.

30. Goude, R., Roberts, D. M. & Parish, T. (2015) Electroporation of mycobacteria, Methods Mol Biol. 1285, 117–30.

31. Schindelin, J., Arganda-Carreras, I., Frise, E., Kaynig, V., Longair, M., Pietzsch, T., Preibisch, S., Rueden, C., Saalfeld, S., Schmid, B., Tinevez, J. Y., White, D. J., Hartenstein, V., Eliceiri, K., Tomancak, P. & Cardona, A. (2012) Fiji: an open-source platform for biological-image analysis, Nat Methods. 9, 676–82.

32. Cangelosi, G. A., Palermo, C. O., Laurent, J. P., Hamlin, A. M. & Brabant, W. H. (1999) Colony morphotypes on Congo red agar segregate along species and drug susceptibility lines in the Mycobacterium avium-intracellulare complex, Microbiology (Reading). 145 (Pt 6), 1317–1324.

33. Etienne, G., Villeneuve, C., Billman-Jacobe, H., Astarie-Dequeker, C., Dupont, M. A. & Daffe, M. (2002) The impact of the absence of glycopeptidolipids on the ultrastructure, cell surface and cell wall properties, and phagocytosis of Mycobacterium smegmatis, Microbiology (Reading). 148, 3089–3100.

34. Santucci, P., Point, V., Poncin, I., Guy, A., Crauste, C., Serveau-Avesque, C., Galano, J. M., Spilling, C. D., Cavalier, J. F. & Canaan, S. (2018) LipG a bifunctional phospholipase/thioesterase involved in mycobacterial envelope remodeling, Biosci Rep. 38.

35. Lee, M. H., Pascopella, L., Jacobs, W. R., Jr. & Hatfull, G. F. (1991) Site-specific integration of mycobacteriophage L5: integration-proficient vectors for Mycobacterium smegmatis, Mycobacterium tuberculosis, and bacille Calmette-Guerin, Proc Natl Acad Sci U S A. 88, 3111–5.

36. Recht, J., Martinez, A., Torello, S. & Kolter, R. (2000) Genetic analysis of sliding motility in Mycobacterium smegmatis, J Bacteriol. 182, 4348–51.

37. Nessar, R., Reyrat, J. M., Davidson, L. B. & Byrd, T. F. (2011) Deletion of the mmpL4b gene in the Mycobacterium abscessus glycopeptidolipid biosynthetic pathway results in loss of surface colonization capability, but enhanced ability to replicate in human macrophages and stimulate their innate immune response, Microbiology (Reading). 157, 1187–1195.

38. Bernut, A., Viljoen, A., Dupont, C., Sapriel, G., Blaise, M., Bouchier, C., Brosch, R., de Chastellier, C., Herrmann, J. L. & Kremer, L. (2016) Insights into the smooth-to-rough transitioning in Mycobacterium bolletii unravels a functional Tyr residue conserved in all mycobacterial MmpL family members, Mol Microbiol. 99, 866–83.

39. Medjahed, H. & Reyrat, J. M. (2009) Construction of Mycobacterium abscessus defined glycopeptidolipid mutants: comparison of genetic tools, Appl Environ Microbiol. 75, 1331–8.

40. Barrow, W. W. & Brennan, P. J. (1982) Isolation in high frequency of rough variants of Mycobacterium intracellulare lacking C-mycoside glycopeptidolipid antigens, J Bacteriol. 150, 381–4.

41. Belisle, J. T., Pascopella, L., Inamine, J. M., Brennan, P. J. & Jacobs, W. R., Jr. (1991) Isolation and expression of a gene cluster responsible for biosynthesis of the glycopeptidolipid antigens of Mycobacterium avium, J Bacteriol. 173, 6991–7.

42. Saviola, B. & Bishai, W. R. (2004) Method to integrate multiple plasmids into the mycobacterial chromosome, Nucleic Acids Res. 32, e11.

